# Evolution of international collaborative research efforts to develop non-Cochrane systematic reviews

**DOI:** 10.1101/467795

**Authors:** I. Viguera-Guerra, Juan Ruano, Macarena Aguilar-Luque, Jesus Gay-Mimbrera, Ana Montilla, J. L. Fernández-Rueda, J. Fernández-Chaichio, J.L. Sanz-Cabanillas, P. Gómez-Arias, Antonio Velez García-Nieto, Francisco Gómez-Garcia, Beatriz Isla-Tejera

**Affiliations:** Agencia de Evaluacion de Tecnologías Sanitarias de Andalucía (AETSA), 14004 Sevilla, Spain; Instituto Maimonides de Investigacion Biomedica de Cordoba (IMIBIC)/Reina Sofia University Hospital/University of Cordoba, Menendez Pidal Ave, 14004 Cordoba, Spain; Department of Dermatology, Reina Sofia University Hospital, Menendez Pidal Ave, 14004 Cordoba, Spain; School of Medicine, University of Cordoba, Menendez Pidal Ave, 14004 Cordoba, Spain; Department of Pharmacy, Reina Sofia University Hospital, Menendez Pidal Ave, 14004 Cordoba, Spain

## Abstract

This research-on-research study describes efforts to develop non-Cochrane systematic reviews (SRs) by analysing demographical and time-course collaborations between international institutions using protocols registered in the International Prospective Register of Systematic Reviews (PROSPERO) or published in scientific journals. We have published an *a priori* protocol to develop this study. Protocols published in scientific journals were searched in MEDLINE/PubMed and Embase databases using the query terms ‘systematic review’[Title] AND ‘protocol’[Title] from February 2011 to December 2017. Protocols registered at PROSPERO during the same period were obtained by web scraping all non-Cochrane records with a Python script. After excluding protocols with less than 90% fulfilled or duplicated, they were classified as published ‘only in PROSPERO’, ‘only in journals’, or in both ‘journals and PROSPERO’. Results of data and metadata extraction using text-mining processes were curated by two reviewers. Datasets and R scripts are freely available to facilitate reproducibility. We obtained 20,814 protocols of non-Cochrane SRs. While ‘unique protocols’ by re-viewers’ institutions from 60 countries were the most frequent, to prepare ‘collaborative protocols’ a median of 6 (2-150) institutions were involved from 130 different countries. Ranked list of countries involved in overall protocol production were the UK, the U.S., Australia, Brazil, China, Canada, the Netherlands, Germany, and Italy. Most protocols were registered only in PROSPERO. However, the number of protocols published in scientific journals (924) or in both PROSPERO and journals (807) has progressively increased over the last three years. *Syst Rev* and *BMJ Open* published more than half of the total protocols. While most productive countries were involved in ‘unique’ and ‘collaborative’ protocols, less productive countries only participated in ‘collaborative’ protocols that were mainly published only in PROSPERO. Our results suggest that although most countries were involved in producing in solitary protocols of non-Cochrane SRs during the study period, it would be desirable to develop new strategies to promote international collaborations, especially with less productive countries.

## Introduction

Systematic reviews (SRs) and meta-analyses (MAs), the standards for evidence synthesis of primary studies, are extremely useful to support decision making processes in the context of Health Systems [1]. However it is desirable that these decisions being supported by reviews of highest methodological quality and have the lowest risk of bias [2]. The Cochrane Handbook for Systematic Reviews of Interventions states that a protocol should be prepared before publishing an SR [3]. In 2010, PRISMA statement advocated registration of SR protocols [1,2]. Preparing an *a priori* protocol will reduce the potential for bias in the review process and increase transparency of analysis and results [4]. Furthemore, if the content of a protocol is available for public access it could also also reduce duplication [5,6], peer review before starting the review process, and audit discrepancies between protocol and the finally produced SR [7–9].

Bias moves away from the possibilities of finding the truth we seek. Rigorously following the methodological agreements established by the scientific community minimizes bias and reduces uncertainty about the estimates we make. One of the proposals to reduce bias in SRs is to develop a comprehensive protocol containing the sources of primary data, procedures for searching, extracting, filtering, selection and analysis of data, as well as the analytical and methodological tools that will be used to conduct the research. For now, it is only essential that such protocols are prepared before making the first analysis and they can be consulted at any time, leaving the trace that were elaborated much before the final results were published [10].

Therefore, there are two main features of a protocol: fist, it should contain all necessary instructions to reproduce the same results from the respective SR; second, it should be prepared before the SR is conducted. Currently, a protocol can be freely viewable to the scientific community with the possibility of tracking dates to make ensure that the protocol is created prior to the review: public repositories and scientific journals. PROSPERO (International prospective register of systematic reviews) is an international database of prospectively registered systematic reviews funded by the National Institute for Health Research (NIHR). Scientists worldwide use the database and less than a decade has reached the landmark achievement of 30,000 registrations. The database is free and open to all researchers planning to conduct an SR and for those searching for registered, ongoing, or completed reviews to develop meta-epidemiology studies. The PROSPERO Advisory Group Statement of Founding Principles centres on free access to both registering and searching the database and is as inclusive as possible, while achieving the key aims of avoiding duplication and minimising bias in SRs.

The second option is to publish the protocol in a scientific journal. This option allows peer reviewers and editors to assess scientifically the quality of the protocol. Through their comments and suggestions, reviewers may suggest changes to improve methodological quality. However, there is still no empirical evidence from meta-epidemiological studies that these changes in protocol will be crucial for improving the quality of the SR. There are many journals that accept protocols of SR for publication. Some of them are ‘BMJ Open’ (https://bmjopen.bmj.com/) and ‘systematic Reviews’ (https://goo.gl/mFShxv).

No matter where protocols are published, a descriptive analysis of these documents can give us a glimpse of the efforts made by researchers to carry out SRs. We recently found (data not published) that only a small percentage of these protocols ends up being published their results, and this could be an unknown source of bias not studied so far, similar to those meant for many randomized clinical trials (RCTs)that were registered in public repositories such as ClinicalTrials.gov but never been published. In a first approximation to this problem, we aimed to describe comprehensively how different reviewers represented by the institutions and countries to which they belong, make efforts to design these protocols and their strategies to conduct reviews.

To date, no study has formally assessed the relationships between reviewers, represented by their respective institution or country, when elaborating an *a priori* protocol to develop a non-Cochrane systematic review, analysing co-working patterns, and evolution of strategies to make these protocols publicly accessible.

## Materials and methods

### *A priori* published protocol

We published an *a priori* protocol in Systematic Reviews [11]. This protocol describe source of data, methods to perform document searching (PROSPERO web scraping and literature databases query), eligibility and screening, data extraction, analysis, and reporting of results.

### Web scraping and literature search strategies

Records stored in PROSPERO (https://www.crd.york.ac.uk/prospero/) were obtained by web scraping using a custom Python 3.0 script and the Chrome’s Web Scraper website data extraction tool (http://webscraper.io/) to automatically and iteratively extract the raw data of all the completed non-Cochrane registration records stored from February 2011 to December 2017. Protocols published in scientific journals were obtained by querying PubMed/MEDLINE and Embase using the *RISmed* R package and the Boolean terms combination ‘[Systematic Reviews”[Title] AND “protocol”[Title]]’.

### Data filtering and eligibility criteria

Registers or protocols with less than 90% of sections fulfilled or those that were duplicated (i.e., those sharing titles and reviewers) were dropped from the dataset. An R script automatically performed the screening process. Subsequently, the results were subjected to human verification by two reviewers (JG-M and MA-L).

### Dataset and variables

A working .csv file, which included only those variables we were interested in for further analysis, was obtained (Supplementary methods). Protocols with different reviewer’s affiliation countries were considered to be the result of international collaboration and their respective countries were analysed as they co-appeared in the protocol as unlisted and tagged as contributing to ‘Collaborative protocols’. Protocols with unique reviewers’ affiliation country were considered to be produced by a unique country and were tagged as ‘Unique protocols’.

### Demographics and evolution of protocol production

Some panels of plots represent changes in the number of protocols published from 2011 (the year PROSPERO was launched) to 2017 (the year PROSPERO web scraping was performed), considering the source (journal vs PROSPERO), type of journal, and country. Considering the entire list of affiliation-associated countries for all co-reviewers per protocol, we displayed a world map that represents in different colours the number of times any country has been involved in any protocol.

### Data visualization, and statistical analysis

Graphs were produced and statistics were analysed using several packages of R 3.4.4 language [R Development Core Team(http://www.R-project.org], except for Venn diagram, obtained using the *eulerr* shinny app (http://eulerr.co), and the workflow figure, created using Review Manager 5 (RevMan 5) software (https://community.cochrane.org/help/tools-and-software/revman-5).

Our analysis can be fully reproduced by using several source files containing raw data and R scripts are available as an R notebook (https://github.com/info4cure/PROSPERO_protocols_demographics). It is shared as open source under the MIT license. A Python script for PROSPERO web scraping will be publically retrieved by the end of 2018, once our team performs all the analyses related to the main project.

### Protocol vs. meta-epidemiological study

Our planned search strategy was published in *Systematic Reviews* and compared with the final reported review methods. The methods of web scraping, filtering and selection did not changed. However, as this is the first article, the project constitutes only a partial descriptive analysis as compared with the main goals described in the above mentioned protocol.

### Ethical considerations

Since our study did not collect primary data, no formal ethical assessment and informed consent are not required.

## Results

### Search results

After scraping 30,000 PROSPERO records, 5,362 documents were excluded as they were not fulfilled and 903 were duplicated versions of other protocols (Fig. 1). After text mining and manually supervising the obtained dataset, 4,364 protocols were not considered because they were lacking in crucial information required for the analyses. By searching bibliographic databases, we obtained 1,732 protocols of SRs published in scientific journals. Only 807 protocols were shared by both PROSPERO and journal datasets (Fig. 2b).

**Fig 1.**
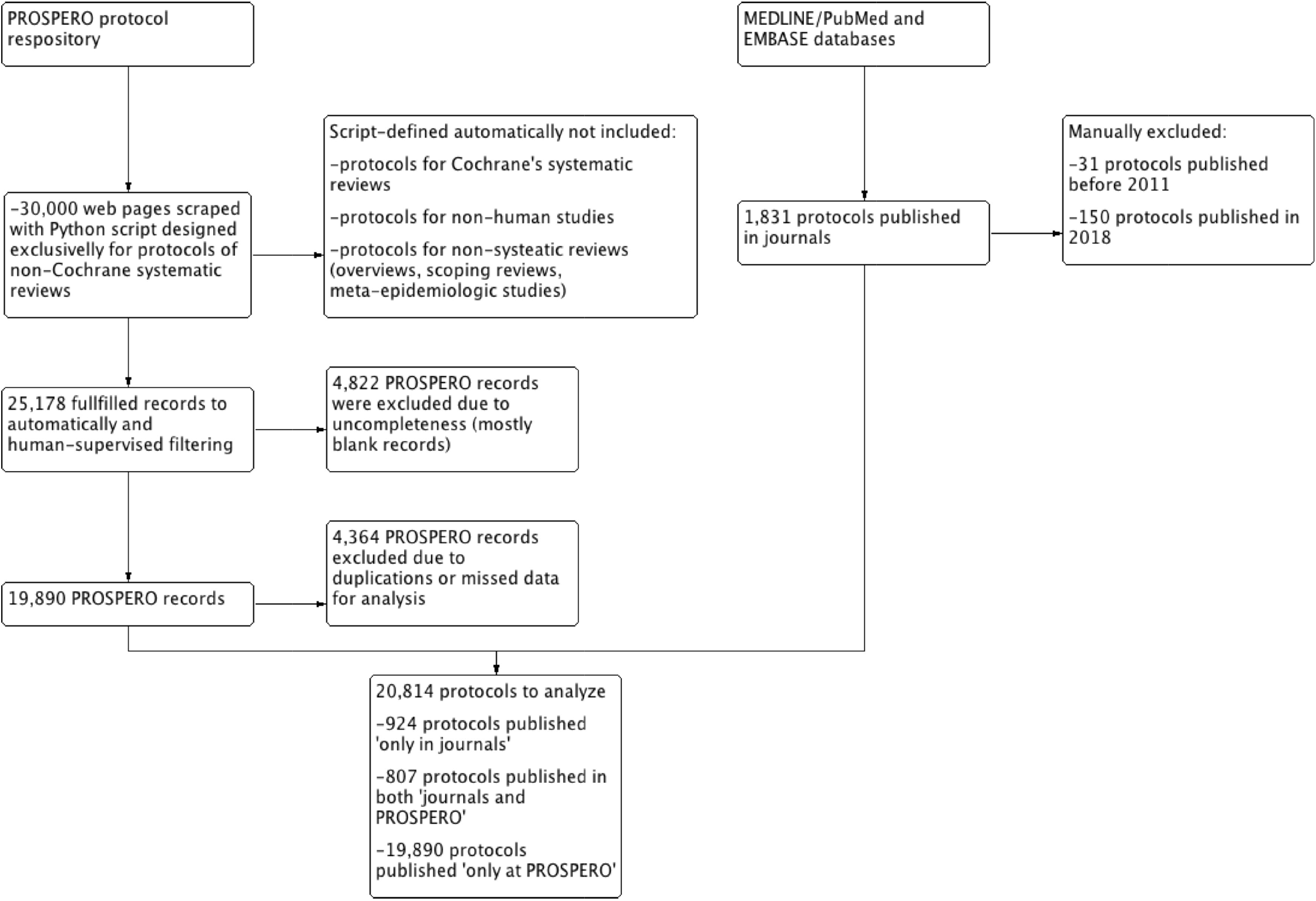
PRISMA workflow of searching for PROSPERO records and protocols published in scientific journals about non-Cochrane systematic reviews.

**Fig 2.**
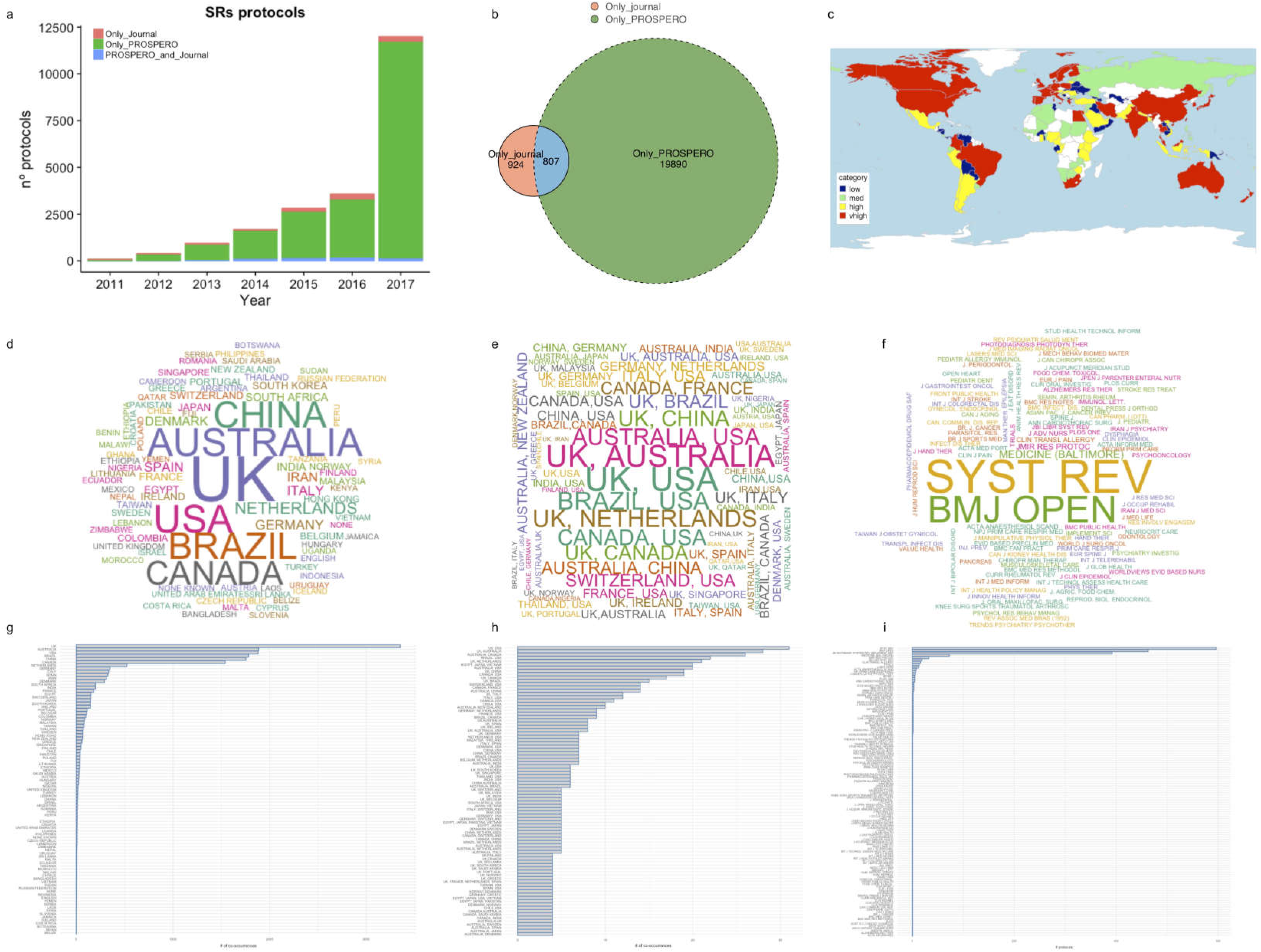
This panel represent the main features of included protocols. (a) Frequency of protocols published from 2011 to 2017 comparing those protocols published ‘only in a journal’, ‘only at PROSPERO’, and in both ‘journal and PROSPERO’. (b) Venn diagram of number protocols published ‘only in a journal’ (coral), ‘only at PROSPERO’ (green), and their intersection, both ‘journal and PROSPERO’ (blue). (c) Map representation of number of protocols produced by country (as proxy of reviewer’s affiliation country). Colours represent levels of productivity defined by quartiles of a new recoded variable [abs(log2(country.count / all.countries.count))] (red, very high; yellow, high; green, medium; blue, low). (d),(e), and (f) represent world clouds of ‘unique countries’ (d), ‘collaborative countries’ (e), and journals (f). Text size and centering is proportional to the associated number of protocols. Colours have been randomly assigned. (g), (h), and (i) represent column plots of ‘unique countries’ (g), ‘collaborative countries’ (h), and journals (i) ranked by total number of protocols.

### General characteristics

Thus, 20,814 protocols published from 2011 to 2017 were finally included for further analyses. These protocols comprised 10,888 reviewers affiliated to institutions across 130 countries. The median number of reviewers and institutions per protocol were five in ranges 1–57 and 1–42, respectively. The total number of produced protocols increased from 2011 to 2017 following and exponential pattern, mainly due to those registered ‘only in PROSPERO’ (Fig. 2a), and followed far behind by protocols published ‘only in journals’ or published in both ‘journal and PROSPERO’ (Fig. 2b).

### Scientific journals vs PROSPERO repository

There were 124 journals where 1,758 protocols of SRs were published (Fig. 3a). Some of these journal by frequency order are ‘BMC Systematic Reviews’ (Syst Rev), ‘British Medical Journal Open’ (BMJ Open), ‘JBI Database of Systematic Reviews and Implementation Reports’, ‘Medicine (Baltimore)’, ‘JMIR Research Protocols’, ‘Clinical and Translational Allergy’, and ‘TRIALS’ (Fig. 2f, 2i, and 3a). More than half of all protocols published in scientific journals may be read in ‘syst Rev’ (33%) or ‘BMJ open’ (27%)(Fig. 3b). The ‘syst Rev’ published maximum protocols during 2011–2017, with the majority (80%) also being registered at PROSPERO (Fig. 3b). This publishing strategy has increased progressively from 2011 onwards, while the number of protocols without PROSPERO registration remained consistently low during the study period (Fig. 3c). This fact seems to be associated with a major number of protocols authored by reviewers affiliated with institutions form the UK and Canada, and to a lesser extent from Australia (Fig. 4c). On the contrary, ‘BMJ Open’, the second journal with more number of published protocols, seems to follow a different pattern of protocol publication. First, the same number of published protocols is available for with vs without PROSPERO registration (45%/55%) for the entire study period (Fig 3b). Second, there is a switch in publication patterns during 2015: published protocols that were also registered in PROSPERO were the majority from the first period (2011–2015), with a peak in protocols from China in 2015 (Fig. 4c). However, since 2016, publishing protocols without being registered in PROSPERO become more frequent, and such protocols were mainly produced by institutions from the UK (Fig 3c).

**Fig 3.**
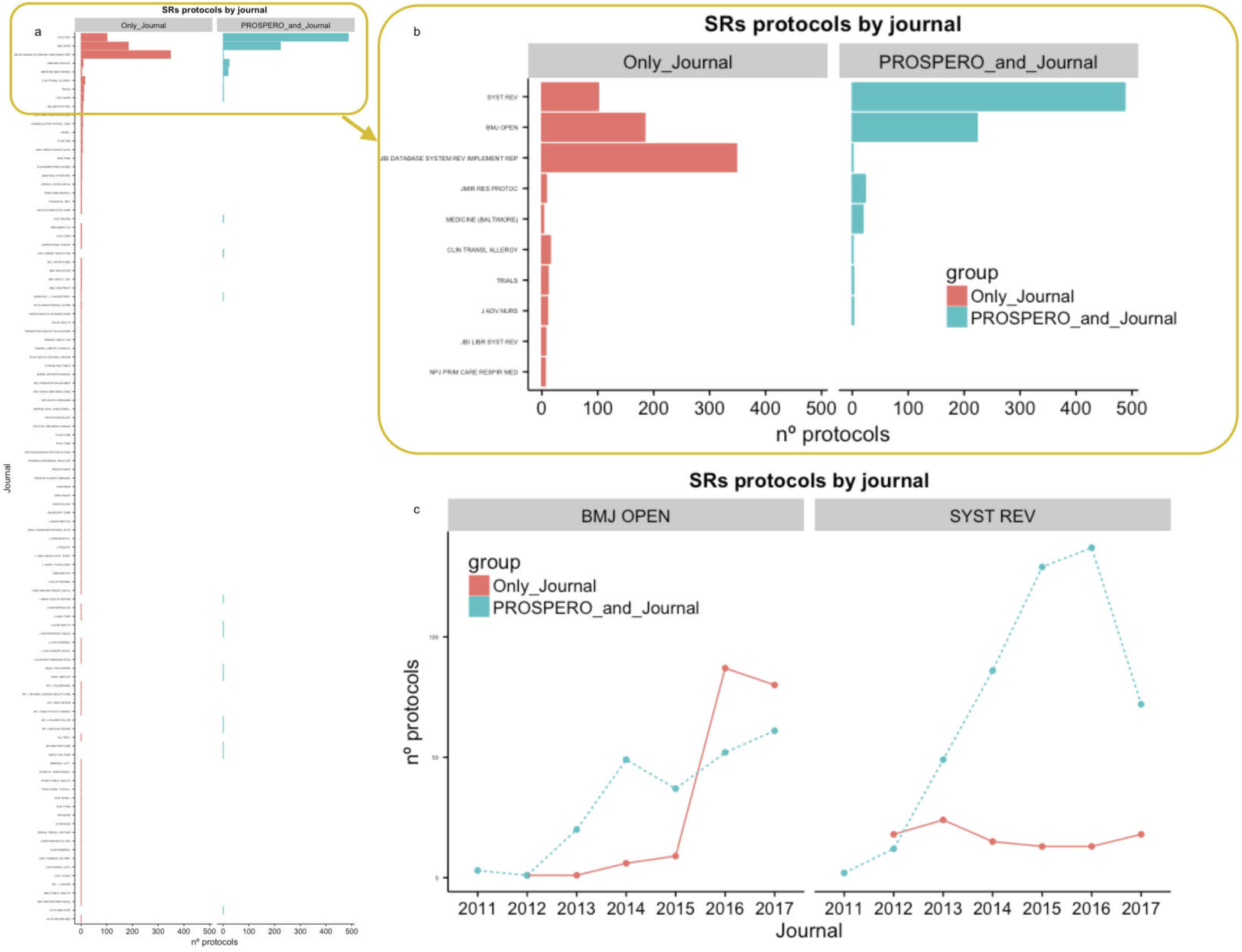
This panel represent frequency and time-course changes of SR protocol publication by journals. (a) Frequency of protocols published from 2011 to 2017 by journal comparing those protocols published ‘only in a journal’ with those protocols published in both ‘journal and PROSPERO’. (b) Magnified vision of plot (a) centered on top 10 most publisher journals. (c) Evolution of ‘only journal’ vs ‘journal and PROSPERO’ protocols publications from 2011 to 2017 comparing ‘BMJ open’ and ‘systematic Reviews’ journals. *SR: Systematic Review*.

**Fig 4.**
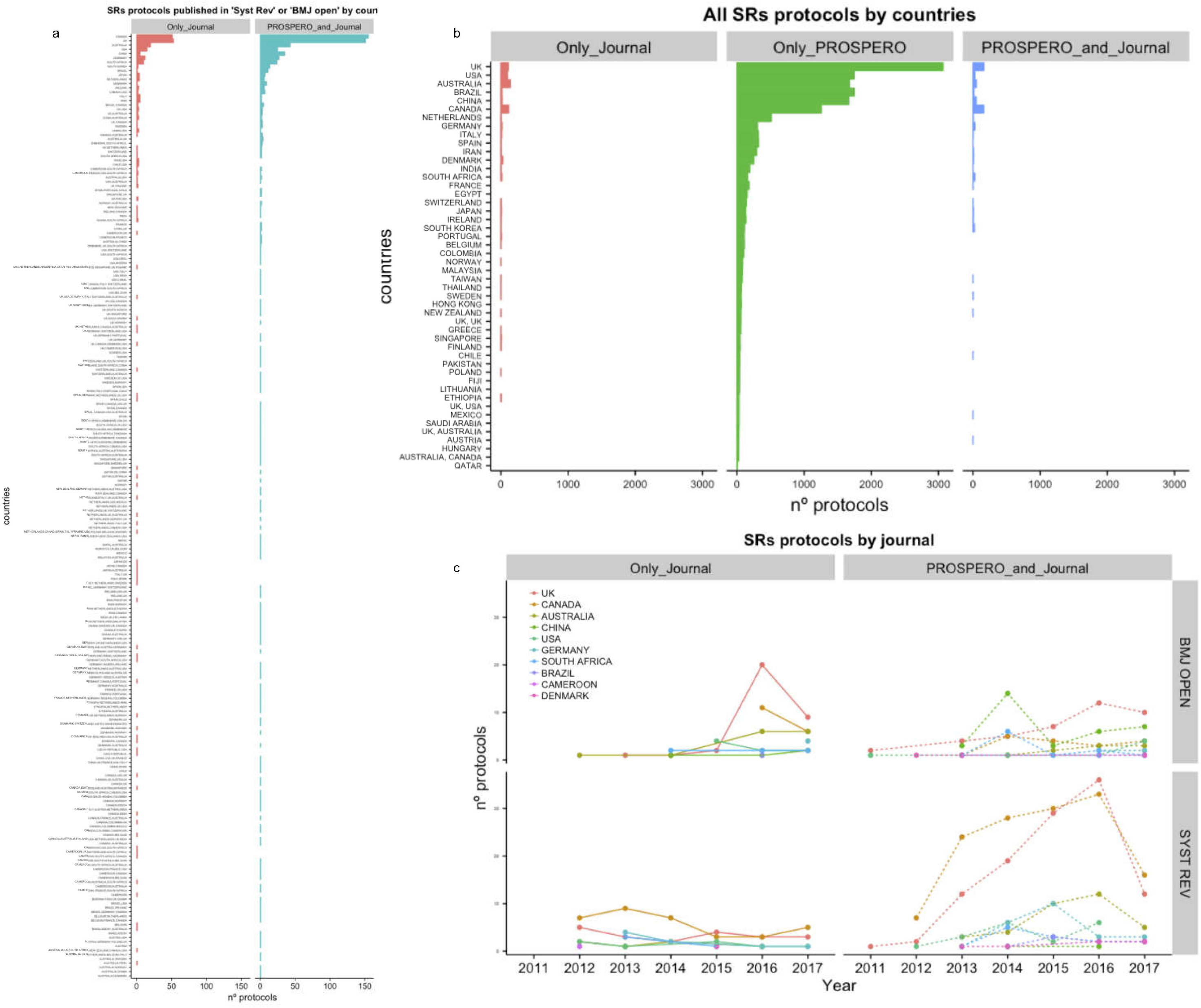
This panel represent frequency and time-course changes of SR protocol publication by countries. (a) Frequency of protocols published from 2011 to 2017 by country comparing those protocols published ‘only in a journal’ with those protocols published in both ‘journal and PROSPERO’. (b) Magnified vision of plot (a) centered on top10 most productive countries comparing those protocols published ‘only in a journal’, “only at PROSPERO’, and in both ‘journal and PROSPERO’. (c) Evolution of ‘only journal’ vs ‘journal and PROSPERO’ protocols publications from 2011 to 2017 comparing top 10 most productive countries.

### Protocols by reviewers’ affiliation countries

Reviewers belonging to institutions from the majority of countries have participated in at least on protocol (Fig. 1c). The distribution levels of a country’s participation in the generation of protocols have been very similar considering the different regions worldwide. However, we show that African countries in comparison with other countries, produce lower number of protocols and the lowest number of protocols are produced by most of these countries. Most protocols (17,431; 90%) were authored by reviewers from institutions of a single country from a total of 90 countries (31.5% countries worldwide). In contrast, a few protocols (1,938) were elaborated by collaboration with institutions from two or more countries from a list of 130 (45.62% of 285 countries worldwide) (Fig. 5). The wordclouds (Fig. 1 d-f) and bar diagrams (Fig. 1g-i) show a predominance participation by institutions from countries such as the UK, the U.S., Australia, Brazil, Canada, China, and the Netherlands in producing protocols either in isolation or in collaboration with others countries (Fig. 1e, 1h, and 4b).

**Fig 5.**
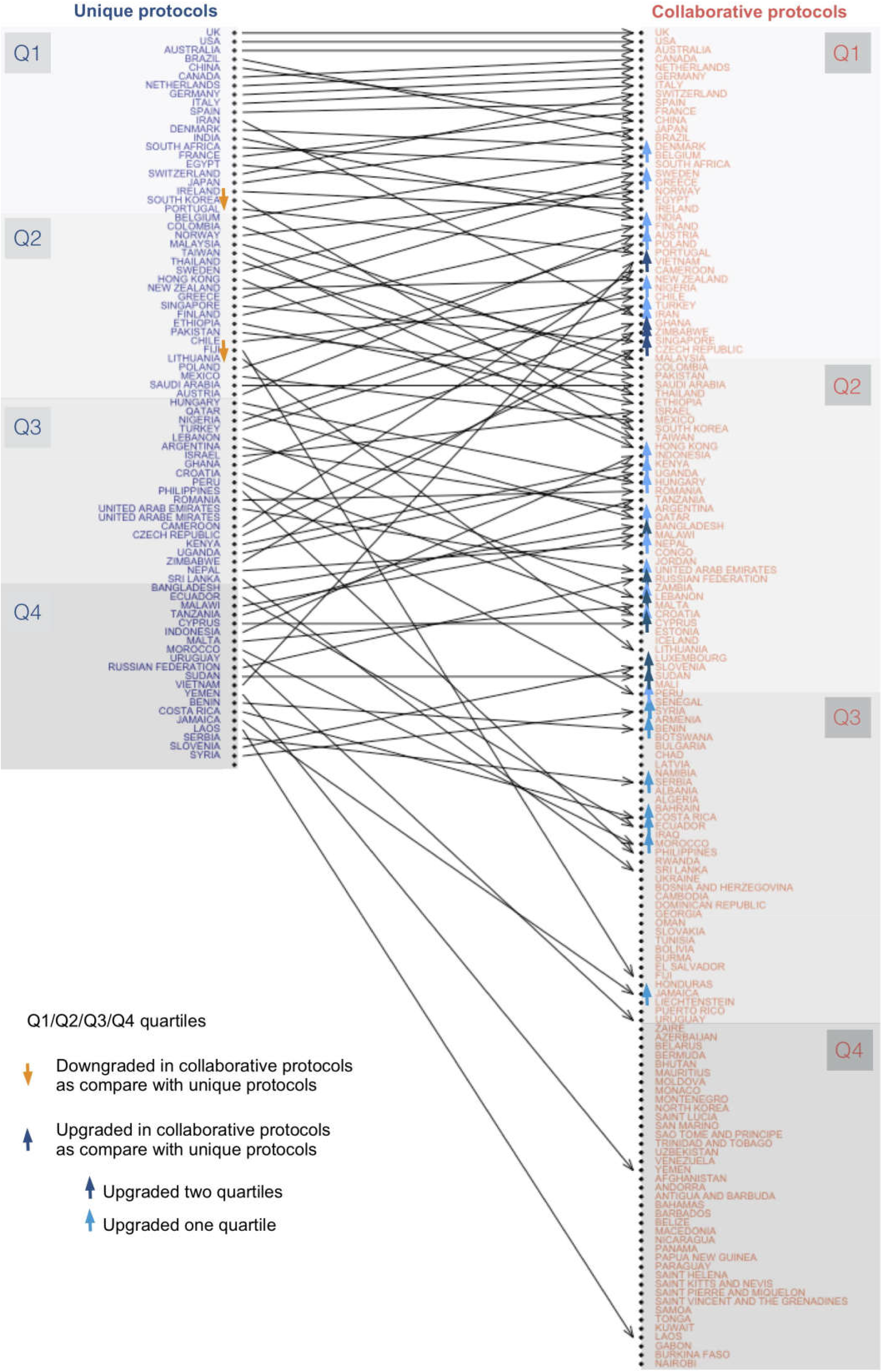
Rank discrepancies between two ordered lists of reviewers’ affiliation countries. The ‘Unique protocols’ column displays a descendent list of reviewers’ affiliation countries that produced protocols for which all reviewers’ institutions belonged to an unique country. The ‘Collaborative protocols’column displays a ranked list of reviewers’ affiliation countries that collaborated with other reviewers’ affiliation countries to produce protocols for SRs. Arrows connect the same country fromfirst to second list. Countries represented only in one of the lists are not connected to/by any arrow. Countries are sub-grouped (Q1:Q4) by cutting through25%, 50%, and 75% of total number of countries in each list. When comparing ‘Unique protocols’ and ‘Collaborative protocols’ lists, country position is considered being modified if the edge connects two different subgroups (i.e.,Q1→· Q3). *Direction of the change defines ‘upgrading’* (*Q*2 → *Q*1, *Q*3 → *Q*1, *Q*4 → *Q*1,*Q*3 → *Q*2, *Q*4 → *Q*3) *or ‘downgrading’ the rank position of any country*(*Q*1 → *Q*2, *Q*1 → *Q*3, *Q*1 → *Q*4, *Q*2 → *Q*3, *Q*2 → *Q*4, *Q*3 → *Q*4).

Production of collaborative protocols has increased during last years. These protocols are being published in PROSPERO since 2011, and later also in journals (2013) or only in journals (2014), respectively (Fig. S1 of Supplementary Information). Protocols from more great number of countries (more than 30) have appeared since 2016, with more than 150 different countries participating in protocols in 2017. The list of countries ranked by participation in collaborative protocols is characterized by: a) being larger than the list of countries producing unique protocols; b) the top 10 countries are repeated in both lists, except that China and Brazil were substituted by the Netherlands and France in the collaborative list; c) almost all the countries involved in producing ‘unique protocols’ participated in the creation of ‘collaborative protocols’ as this is more productive than working in isolation; d) countries that only participated in collaborative protocols were the least productive (Fig. 5).

### Analysis of time-course patterns

Fig. 6 displays ‘the year of the first protocol published’ by ‘source of publication (‘only PROSPERO’, ‘PROSPERO journal’, ‘only journal’)’ and by ‘country’. To simplify the analysis, only protocols without collaborations between countries are considered in this plot. There seem to be four different patterns. In the first pattern, the most productive and collaborative countries (the UK, Canada, Australia, China, Germany, Italy, the U.S., and the Netherlands) started producing protocols very early and submitted them to PROSPERO or published in journals; however, it was only after 1-3 years that they started publishing protocols in both PROSPERO and journals. In the second pattern, countries (Denmark, Brazil, Ireland, South Africa, Spain, India, Taiwan, Iran, New Zealand, Japan, Switzerland, Sweden, South Korea, and France) started publishing their protocols in PROSPERO and, after 1-3 years in PROSPERO and journals, and finally, after another 1-4 years, they finally started publishing their protocols only in journals. The third pattern defines countries (from Belgium to United Arab Emirates) that started using only PROSPERO, not too early, and most of them, after a longer period of 3-5 years, published only in journals without submitting their protocols to PROSPERO anytime. In the final pattern, least productive countries covering from Hong Kong to Namibia, and representing more than half of the world’s countries, submitted their protocols only to PROSPERO.

**Fig 6.**
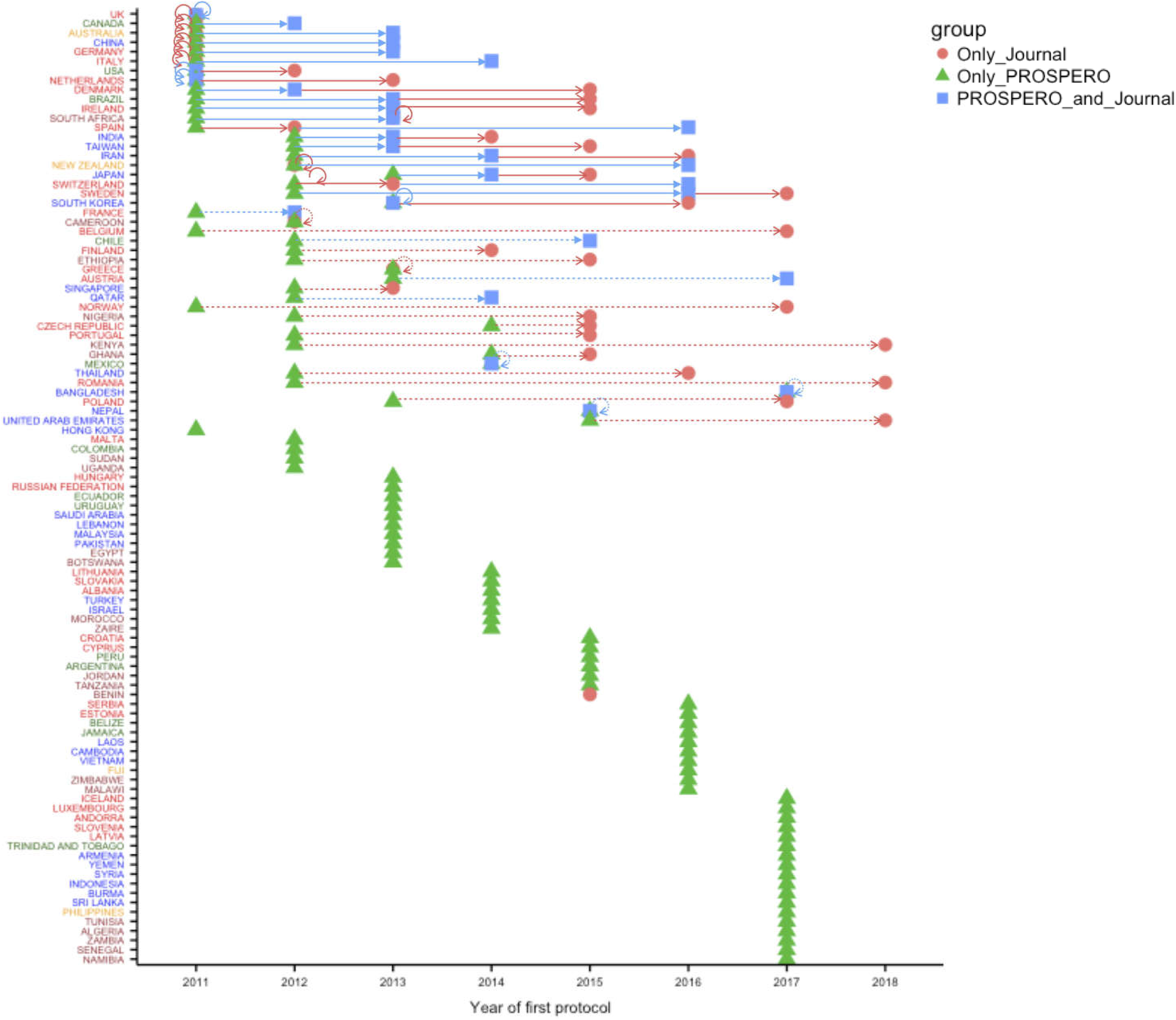
Analysis of protocol publication patterns by most productive countries. Countries are listed in a descendent order based on their ‘Unique protocol’ productivity. Points represent a hallmark in every country’s history of protocol publication: first time to publish a protocol in ‘only a journal’ (red dot), ‘only at PROSPERO’ (green triangle), and in both ‘journal and PROSPERO’ (blue square). Arrows connect two (by a dotted line) or more (by a full line) hallmarks to emphasize how much time is taken for a country to adopt a new way of publication.

## Discussion

### Main findings

This is the first study describing the diversity of collaborative strategies and time-course preference changes followed by countries whose institutions involved in producing and communicating protocols for SRs. Overall, our findings suggest three observations: first, most countries are involved in producing protocols for SRs, although the majority of protocols are produced without international collaboration; second, although most of protocols were earlier registered only in PROSPERO, this tendency seems to be changing since 2013-2014 and most productive countries have begun publishing protocols mainly in ‘syst Rev’ or ‘BMJ Open’journals; third, less productive countries participate through international collaborations to conduct protocols that are predominantly submitted to PROSPERO and not to scientific journals.

### Our findings in context

Our results show that most protocols for non-Cochrane SRs were authored by reviewers from institutions belonging to a single country. AS most of the topics are not restricted by local, ethnic, or geographic factors, this approach may become a challenge for transforming evidence to practice worldwide. Collaborations between countries, especially more vs less productive institutions, may enhance technical expertise by training in the later regions and extending collaboration beyond SRs, improving the adoption of evidence-based health policies, selection of the best evidence for the right audience, and focusing on relevant issues through appropriate methodologies [12]. This will be possible with the growing innovation in tools and platforms that would enable more efficient SR production in collaboration.

Registration forms of protocols submitted to PROSPERO are only checked against the scope for inclusion in the repository and for clarity of content. Once accepted an audit trail of major changes to planned methods may be checked at any time, even after the SR published. Ideally, registration should take place before the researchers started formal screening against inclusion criteria, but reviews are eligible as long as they have not progressed beyond the point of completing data extraction. However, accomplishing these goals is still conditioned by the reviewers integrity. Dawid Pieper and Katharina Allers recently suggested than *a priori* design of an SR may not have the same advantages and potential to reduce the risk of bias in SRs compared to those of RCTs. They argue that future ‘living SRs’, involving new workflow and collaboration tools, text mining and machine learning technologies, emerging reusable data depositories, and shared ontologies and harmonized data transfer protocols can be expected to decrease any potential manipulations in future.

Since 2011, there have been few meta-epidemiological studies about the content and use of PROSPERO. A descriptive analysis of the number of PROSPERO registrations and website usage from 2011 to 2017 have recently published, exploring the epidemiological characteristics and completeness of primary outcome pre-specification in a small sample of PROSPERO records [13]. They highlight the exponential increase in registered protocols at PROSPERO between 2011 and 2017. However, these authors recognize that there are still many caveats regarding the real utility of making a document, which is not methodologically reviewed, available of public access. These authors, one of them a member of the PROSPERO’s international advisory group, raise three issues that will certainly generate future debate about new strategies for future improvement in PROSPERO: how closely published SRs adhere to the planned methods-PROSPERO registrations?; can specification of greater outcomes in PROSPERO registrations prevent inclusion and reporting biases?; and do registered SRs address the necessary questions?.

### Limitations and strengths

Our study includes analyzing the largest sample of SR protocols produced during the last seven years. We did not perform PROSPERO registration record sampling [13]. Rather, our objective was to get the entire universe of registers, from the first document to the last one registered just before the date of web scraping, and not a representative sample of them. The search specificity for non-Cochrane PROSPERO registration records was based on Python script that was designed to recognize only the format of these records, which differs from registration records for Cochrane and non-human studies. These cannot be scraped using our script due to the structural differences in PROSPERO forms. Thus, the sensitivity and specificity for the web scraping is 100%.

The present study, however, also had several limitations. First, we used countries as a proxy for reviewers or reviewers’ institutions. Better size of information granularity would have enable deeper analysis about reviewers’ and institutions’ productivity and collaborative networks. However, we decided based on technical issues related to the variety of formats used by reviewers when fulfilling PROSPERO forms and time limitation to afford the project to use countries as the unit of analysis. We have used this approach previously to analyse how the author-paper affiliation network architecture influences the methodological quality of SRs and MAs of psoriasis [14]. Second, our study is limited to protocols in non-Cochrane SRs. Cochrane Reviews are demonstrated to have better methodological quality and lower bias risk than non-Cochrane SRs [15,16]. The Cochrane Database of Systematic Reviews (CDSR) is the leading resource for Cochrane SRs and protocols. Protocols for Cochrane Reviews have also been published at PROSPERO from 1 October 2013. This fact introduces a gap at PROSPERO from 2011 to 2013 with only non-Cochrane SRs. Another area of particular concern in relation to non-Cochrane reviews is the failure to register reviews at the outset. Registration of Cochrane reviews is mandatory with publication of a protocol *a priori*. To avoid an unbalanced sample of protocols with different proportions of quality and rate of publication, we have decided to select only non-Cochrane SRs protocols for our study.

### Implications of results

Future work should focus on analyzing co-authorship networks. This would help to identify academic talent, put the head-to-head research interest of experienced researchers and local investigators who have recognised regional resources and time changing health-care necessities, thus increasing the opportunities for improving international collaboration. A recent article has demonstrated, after mapping of 115,000 RCTs, the mismatch between research efforts and health needs in non–high-income regions [17]. Similarly, implementing strategies to efficiently coordinate collaborations between countries to perform non-Cochrane SRs, and this is especially so when most human resources and stakeholders are not coordinated by a consolidated international organization such as the Cochrane.

Our results demonstrate that most protocols are registered only in PROSPERO. However, before reviewers beguin developing SRs, PROSPERO registrations can not be methodologically curated, critically per-reviewed, or freely commented on by anonymous readers. If protocols should be *a priori* submitted to a public repository and/or to a journal is debatable. Indeed, our data show that this tendency seems to be changing since 2013–2014 and most productive countries have begun publishing protocols in ‘syst Rev’ or ‘BMJ Open’ journals.

Future studies comparing methodological quality of protocols registered ‘only in PROSPERO, ‘only in journals’, and in ‘both PROSPERO and journals’ should provide empirical evidence for an *a priori* peer-reviewing process of protocols before authors start SR development. Furthermore, by exploring if modifications suggested by peer-reviewers after submitting an SR protocol to a journal significantly improves the quality of protocols (i.e. assessed using PRISMA for Protocols extension), and even to increase the methodological quality and to reduce the risk of bias in the final SR would be of great interest.

### Conclusions and Future research

Although most countries worldwide were involved in producing protocols for non-Cochrane SRs, it is desirable to develop new strategies to boost international collaborations, especially between more productive and less productive countries. While most protocols of SRs are submitted to PROSPERO, the potential advantage of a new tendency to publish protocols in scientific journals and not only in PROSPERO should be evaluated in future.

## Supporting information

## Acknowledgments

We wish to acknowledge the assistance received from Dr. Dawid Pieper (Head of Department, Evidence-based health services research, Institute for Research in Operative Medicine, Faculty of Health, School of Medicine at Witten/Herdecke University, Cologne, Germany) for answering our questions of interpreting patterns of collaborative relationship between countries during the analysis phase of our study. We are grateful to all the contributors and to the advisory group of the PROSPERO registry. We are also grateful to the reviewers for their helpful comments and suggestions. We would also like to thank Editage (www.editage.com) for English language editing.

## Author contribution

Conceptualization: IV-G and JR. Funding acquisition: JR. Project administration: IV-G and JR. Resources: IV-G and AM. Data scraping: JLF-R and JF-Ch. Data curation: MA-L, JG-M, and AM. Formal Analysis: IV-G, JR. Visualization: IV-G and JR. Methodology: IV-G, JLS-C, and PG-A. Supervision: AVG-N and BI-T. Writing - original draft: IV-G and JR. Writing - review and editing: JR, AVG-N, and BI-T.

## Supporting Information

### S1 File

**1. Supplementary methods. 1.1. Web scraping and literature search strategies. 1.2. Dataset and variables**

## References

1. Abuabara K, Freeman EE, Dellavalle R. The role of systematic reviews and meta-analysis in dermatology. J Invest Dermatol 2012;132:e2.

2. F. Gomez-Garcia, J Ruano, J. Gay-Mimbrera, M. Aguilar-Luque, J.L. Sanz-Cabanillas, P. Alcalde-Mellado, et al. Most systematic reviews of high methodological quality on psoriasis interventions are classified as high risk of bias using ROBIS tool. J Clin Epidemiol 2017;92:79–88.

3. J.P.T. Higgins, S. Green (Eds.), Cochrane handbook for systematic reviews of interventions, Version 5.1.0 [updated March 2011], The Cochrane Collaboration 2011.

4. D Moher, L Shamseer, M Clarke, D Ghersi, A Liberati, M Petticrew, et al. Preferred reporting items for systematic review and meta-analysis protocols (PRISMA-P) 2015 statement. Syst Rev 2015;4:1.

5. K.C Siontis, T. Hernandez-Boussard, J.P.A. Ioannidis. Overlapping meta-analyses on the same topic: survey of published studies. BMJ 2013;347:f4501.

6. Ioannidis JP. The Mass Production of Redundant, Misleading, and Conflicted Systematic Reviews and Meta-analyses. Milbank Q 2016;94:485–514.

7. Kirkham JJ, Altman DG, Williamson PR. Bias due to changes in specified outcomes during the systematic review process. PLoS One 2010;5:e9810.

8. Pandis N, Fleming PS, Worthington H, Dwan K, Salanti G. Discrepancies in Outcome Reporting Exist Between Protocols and Published Oral Health Cochrane Systematic Reviews. PLoS One 2015;10:e0137667.

9. Tricco AC, Cogo E, Page MJ, Polisena J, Booth A, Dwan K, MacDonald H, Clifford TJ, Stewart LA, Straus SE, Moher D. A third of systematic reviews changed or did not specify the primary outcome: a PROSPERO register study. Journal of Clin Epidemiol 2016;79:46–54.

10. Moher D, Shamseer L, Clarke M, Ghersi D, Liberati A, Petticrew M, Shekelle P, Stewart LA. Preferred Reporting Items for Systematic Review and Meta-Analysis Protocols (PRISMA-P) 2015 statement. Syst Rev 2015;4:1.

11. Ruano J, Gómez-García F, Gay-Mimbrera J, et al. Evaluating characteristics of PROSPERO records as predictors of eventual publication of non-Cochrane systematic reviews: a meta-epidemiological study protocol. Syst Rev 2018;7:43.

12. Su TT, Bulgiba AM, Sampatanukul P, Sastroasmoro S, Chang P, Tharyan P, et al. Clinical Epidemiology (CE) and Evidence-Based Medicine (EBM) in the Asia Pacific region (Round Table Forum). Prev Med 2013;57 Suppl:S5–7.

13. Page MJ, Shamseer L, and Tricco AC. Registration of systematic reviews in PROSPERO: 30,000 records and counting. Syst Rev 2018;7:32.

14. Sanz-Cabanillas JL, Ruano J, Gomez-Garcia F, Alcalde-Mellado P, Gay-Mimbrera J, Aguilar-Luque M, et al. Author-paper affiliation network architecture influences the methodological quality of systematic reviews and meta-analyses of psoriasis PLoS One 2017;4: e0175419.

15. Fleming PS, Seehra J, Polychronopoulou A, Fedorowicz Z, Pandis N. Cochrane and non-Cochrane systematic reviews in leading orthodontic journals: a quality paradigm? European Journal of Orthodontics 2013;2:244–248.

16. Goldkuhle M, Narayan VM, Weigl A, et al. A systematic assessment of Cochrane reviews and systematic reviews published in high-impact medical journals related to cancer. BMJ Open 2018;8:e020869.

17. Atal I, Trinquart L, Ravaud P, and Porcher R. A mapping of 115,000 randomized trials revealed a mismatch between research effort and health needs in non-high-income regions. Journal of Clinical Epidemiology 2018;98;123–32.

