## Supplementary material for "Evolution of international collaborative research efforts to develop non-Cochrane systematic reviews"

[a] Agencia de Evaluación de Tecnologías Sanitarias de Andalucía (AETSA), 14004 Sevilla, Spain
[b] IMIBIC/Reina Sofía University Hospital/University of Cordoba, 14004 Córdoba, Spain
[c] Department of Dermatology, Reina Sofía University Hospital, 14004 Córdoba, Spain
[d] Department of Pharmacy, Reina Sof´ıa University Hospital, 14004 Córdoba, Spain
[e] School of Medicine, University of Cordoba, 14004 Córdoba, Spain

### 1. Supplementary methods

*1.1. Web scraping and literature search strategies*
We ran the custom script from 25 November to 1 December, 2017. The script iterated through web pages by modifying URL with for loop changing the RecordID value by choosing a number from 00000 to 80000\footnote{https://www.crd.york.ac.uk/prospero/display_record.php? RecordID=XXXXX}. All obtained data were stored locally as .csv files, where rows represented protocols and columns protocol sections. All PROSPERO records and published protocols were tabulated in a .csv file,

*1.2. Dataset and variables*
*Variables we were interested in for further analysis: a) Reviewers/authors: 'reviewer/author affiliation's country'; b) Articles: 'journal name', 'year of publication'; c) PROSPERO records: 'PROSPERO ID', 'year of registration'.  After performing some in/out joint matches by using PROSPERO ID between records in PROSPERO and protocols published in journals, we classified every unit of analysis as published 'only in journal', 'only in PROSPERO', or in both 'journal and PROSPERO'.*
